# Thermodynamic profiles for co-translational trigger factor function

**DOI:** 10.1101/2023.08.23.554456

**Authors:** Therese W. Herling, Anaïs M. E. Cassaignau, Anne S. Wentink, Quentin A. E. Peter, Pavan C. Kumar, Tadas Kartanas, Matthias M. Schneider, Lisa D. Cabrita, John Christodoulou, Tuomas P. J. Knowles

## Abstract

Molecular chaperones are central to the maintenance of proteostasis in living cells. A key member of this protein family is trigger factor (TF), which acts throughout the protein lifecycle and has a ubiquitous role as the first chaperone encountered by proteins during synthesis. However, our understanding of how TF achieves favourable interactions with such a diverse substrate base remains limited. Here, we use microfluidics to reveal the thermodynamic determinants of this process. We find that TF binding to empty 70S ribosomes is enthalpy-driven, with micromolar affinity, while nanomolar affinity is achieved through a favourable entropic contribution for both intrinsically disordered and folding competent nascent chains. These findings suggest a general mechanism for co-translational TF function, which relies on occupation of the exposed TF substrate-binding groove, rather than specific complementarity between chaperone and RNC. These insights add to our wider understanding of how proteins can achieve broad substrate specificity.

---

Biological function is underpinned by non-covalent and transient protein interactions, which rely on structure and dynamics to achieve selectivity and specificity in the crowded environment of the cell. Molecular chaperones in particular, have evolved towards such interactions, supporting protein folding and prevent misfolding, and key chaperones such as TF act on a remarkably diverse range of substrates.^1–11^ In bacteria, TF is the first chaperone encountered by the nascent polypeptide emerging from the ribosome during synthesis.^1–3, 12–15^ To support proper cellular function, TF operates in a network of non-covalent interactions targeting a broad range of unfolded client proteins within the crowded environment of the cytosol, Figure 1**a**.^3, 6, 16, 17^ Here, we investigate whether TF-ligand binding for a diverse set of substrates shares a common free energy profile.

**Figure 1:**
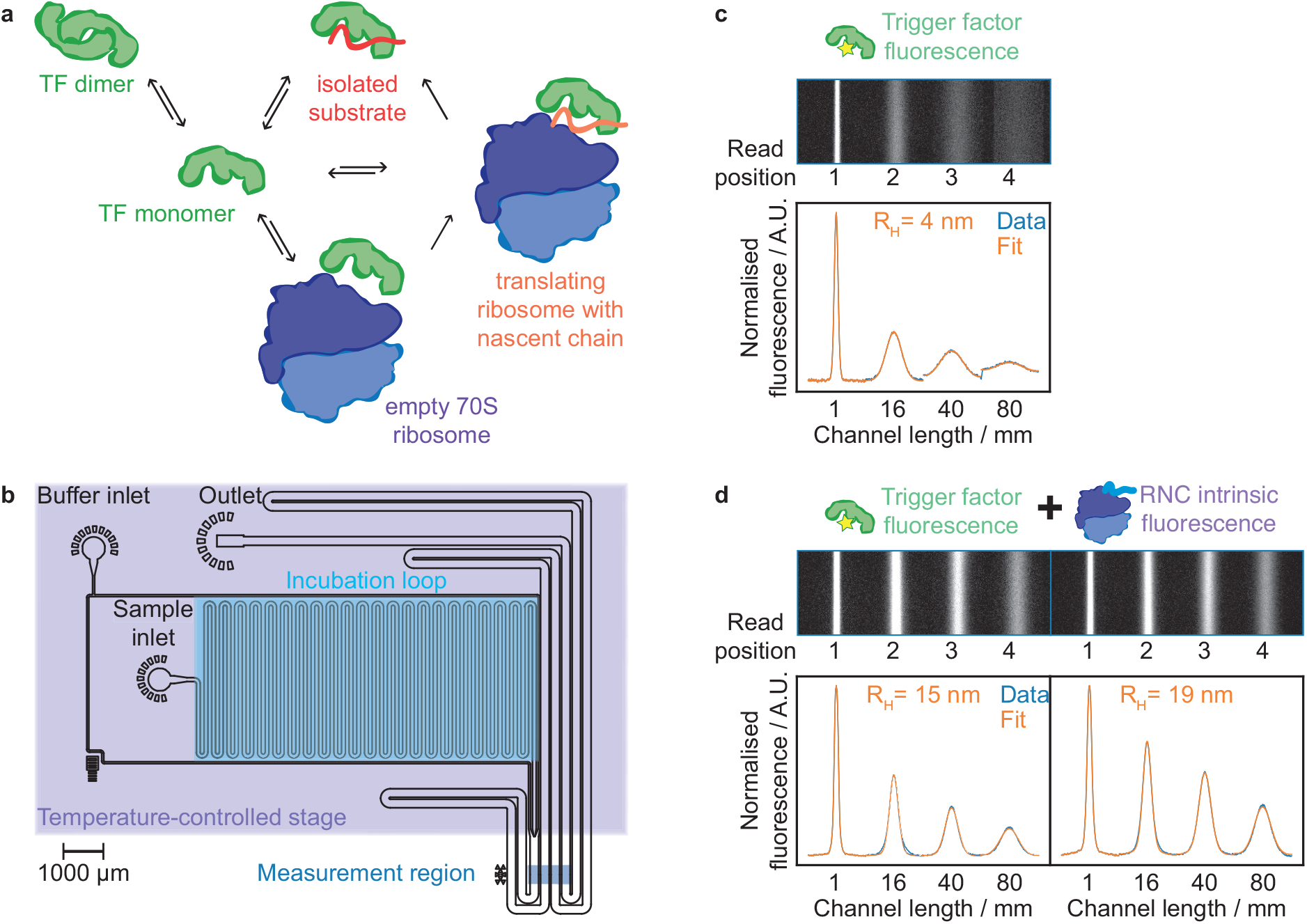
Methodology.**a** The network of TF interactions includes binding to isolated proteins, empty ribosomes, RNCs and dimerisation. **b** Microfluidic diffusional sizing enables the hydrodynamic radius of biomolecules to be determined in free solution. The microfluidic chip is used in conjunction with a temperature-controlled stage to characterise the thermodynamics of protein interactions. **c** Fluorescence image of 200 nM AlexaFluor488 labelled trigger factor in the measurement region of the diffusional sizing chip. Below, the corresponding fluorescence profiles in blue, with a fit to the data in orange in order to obtain *D* and *R*_*H*_ . **d** Analysis of multiple components in a mixture 200 nM trigger factor (left) and the intrinsic fluorescence from 4 *μ*M luciferase RNC (right).^34^ Binding to the RNC is measured through the increase in TF *R*_*H*_.

The importance of TF in cellular function and malfunction has resulted in considerable research on the molecular mechanisms behind the function of this chaperone.^1, 2, 8, 9, 13, 14, 16–23^ New roles for TF are emerging, including in protein secretion and degradation pathways.^8, 24, 25^ These functions are in addition to the action of TF as a general co-translational chaperone,^1, 3, 4, 7, 8, 13, 16, 19^ anti-aggregation chaperone,^17, 23^ unfoldase,^9^, and peptidy-prolyl cis/trans isomerase.^22, 26^ Indeed, the chaperone actively changes the conformational search of its substrates,^10^ it can promote folding against an applied force and modulates the pulling force on nascent polypeptides during translation.^15, 21^ In addition, trigger factor cooperates with bacterial release factor 3 to terminate misfolded nascent chains.^11^

Oligomerisation is a common trait for many molecular chaperones,^27^ and TF self-associates to form a dimer with an equilibrium dissociation constant (*K*_*d*_) typically found to be 1-2 *μ*M,^3, 4, 13^ although *K*_*d*_ values as high as 18 *μ*M have been reported,^16^ Figure 1**a**. TF has an elongated structure, and dimerisation buries the large substrate-binding groove, leading to only a small increase in the observed radius.^28, 29^ The rates for dimer dissociation and association are high (i.e. 6 *·* 10^6^ *M* ^−1^ *s*^−1^ and 10 *s*^−1^)^26^ compared to those for binding to client proteins (typically 10^4^ − 10^5^ M^−1^ s^−1^ and 0.05 s^−1^).^4^ The dimer is therefore considered as a storage unit for the chaperone, which can readily be mobilised to meet substrate demand.^3, 27^

TF functions in a complex network of interactions with different substrate types ranging from small isolated proteins to MDa RNCs, Figure 1**a**, and a multidisciplinary research effort has focussed on elucidating the structural, equilibrium and kinetic parameters for TF function.^1, 2, 8, 9, 13, 14, 16–23^ Nuclear magnetic resonance (NMR) has provided structural and dynamic insight into interactions with misfolded proteins,^17^ RNCs,^19^ and dimer formation.^26, 29^ Fluorescence-based methods have been particularly useful in providing information on the dynamics between TF and actively translating ribosomes.^3, 4, 7^ These studies have shown that TF associates with the RNC at the exit tunnel,^7, 13, 14^ and that the chaperone can detach from the ribosome to remain associated with the emerging nascent chain (NC) with a substrate-dependent *t*_1*/*2_ of up to 35-111 s, whereas binding to the ribosome/RNC surface occurs with nanomolar affinity and a *t*_1*/*2_ of ∼10 s.^3, 7^ A proteome-wide *in vivo* study showed weak TF-RNC affinity for nascent chains ≤100 amino acids,^8^ whereas particularly tight binding (2-110 nM) and fast kinetics (t_1_*/*2 = 0.06-1.7 s) were reported for 75 amino acid NCs.^18^ Despite elegant structural, equilibrium, and kinetic investigations of TF function, it remains poorly understood how the chaperone achieves high affinity for diverse nascent chain sequences.

In particular, the free energy contributions that drive TF-RNC interactions have been challenging to access. The main challenges in probing these systems are the large size range of the interaction partners involved (kDa to MDa), the wide range of interaction affinities (nM to *μ*M), and the need to work across a range of temperatures. Specific probes (e.g. optical or magnetic) can have very high sensitivity, but typically only perform optimally in a section of the required parameter space. To cover a wide range of molecular weights, affinities and temperature ranges, we focus on measurements of a fundamental property, the physical size, of the molecular components as they interact. We measure size (hydrodynamic radius, *R*_*H*_) through monitoring changes in the molecular diffusion coefficients and electrophoretic mobilities (*μ*_*e*_) of the molecular components confined in microfluidic channel that provide highly stable flow conditions with no convective mixing, Figure 1**a-d** and SI Figure 1.^30, 31^ The microfluidic assays can accommodate a wide range of sample dimensions, and we have used them to characterise samples ranging from small molecules to amyloid fibrils.^30–33^ We analyse multiple components in a mixture by combining intrinsic protein fluorescence from unlabelled RNCs with selective fluorophore-labelling of TF, SI Figure 2,^32, 34^ and in this study, we include a Peltier stage to heat and cool samples on chip.^35^ Taken together, the microfluidic setup offers a general-purpose platform for the study of otherwise challenging systems such as TF.

Using this approach, we determine the *K*_*d*_ for TF-substrate binding as a function of temperature, we create snapshots of co-translational TF interactions with RNCs that have been arrested mid-synthesis by a SecM sequence.^19^ We find that TF binds to RNCs for both intrinsically disordered proteins (IDPs) based on *α*-synuclein (*α*syn) and folding-competent (firefly luciferase) with nanomolar affinity, whilst binding to the empty 70S ribosome occurs with low micromolar affinity. By analysing the change in entropy (ΔS) and enthalpy (ΔH) of substrate binding we discover that all the RNCs investigated here share a general thermodynamic profile where binding is promoted by a positive overall ΔS. This profile is distinct from the enthalpy-driven binding to empty ribosomes. Taken together, our data suggests a model for co-translational TF binding, that relies on favourable entropy from the NC occupying the substrate binding groove of ribosome-bound TF, rather than specific complementarity between the chaperone and NC sequence.

## Results

TF is an ATP-independent chaperone, and the manner by which it achieves a favourable Gibbs free energy (ΔG) for binding to a wide range of substrates is therefore of particular interest.^27^ Information on the entropy and enthalpy for TF-substrate interactions has been challenging to obtain due to the high molecular weight of the functional complexes (*>*2.4 MDa) in combination with the low sample concentrations required to access *K*_*d*_*s* in the nanomolar range, e.g. for TF-RNC association.^3, 7^ In this study, we introduce a temperature-controlled microfluidic setup, which enables us to acquire data in a consistent manner across a range of temperatures and gain insight into the thermodynamic driving forces that promote TF-substrate binding, Figure 1**b** and SI Figure 2**a**.

Exploration of the co-translational role of TF requires an understanding of its dimerisation as well as interactions with isolated proteins, nascent chain complexes and empty ribosomes.^2–4, 7, 17, 19, 26, 29^ Does TF employ a general strategy to achieve a favourable ΔG for substrate binding? To address this question, we created snapshots of TF function by determining the *K*_*d*_ for binding to empty 70S ribosomes and three distinct RNCs (i) a folding-competent firefly luciferase RNC, expressed using the TF knockout strain, *Escherichia coli* Δtig,^36^ (ii) the intrinsically disordered protein *α*-synuclein, and (iii) a chimeric *α*-synuclein(Luc) where residues 87-100 have been substituted for the firefly luciferase sequence, a hydrophobic motif characteristic of strong TF binders, Figure 2**a**.^3, 4, 7, 9, 19^ Initially, we determined the *K*_*d*_ for TF dimerisation using free-flow electrophoresis and obtained a value of 1.5 *±* 0.25 *μ*M, SI Figure 1. A TF concentration of 200 nM was therefore chosen for our study to ensure a predominantly monomeric TF.

**Figure 2:**
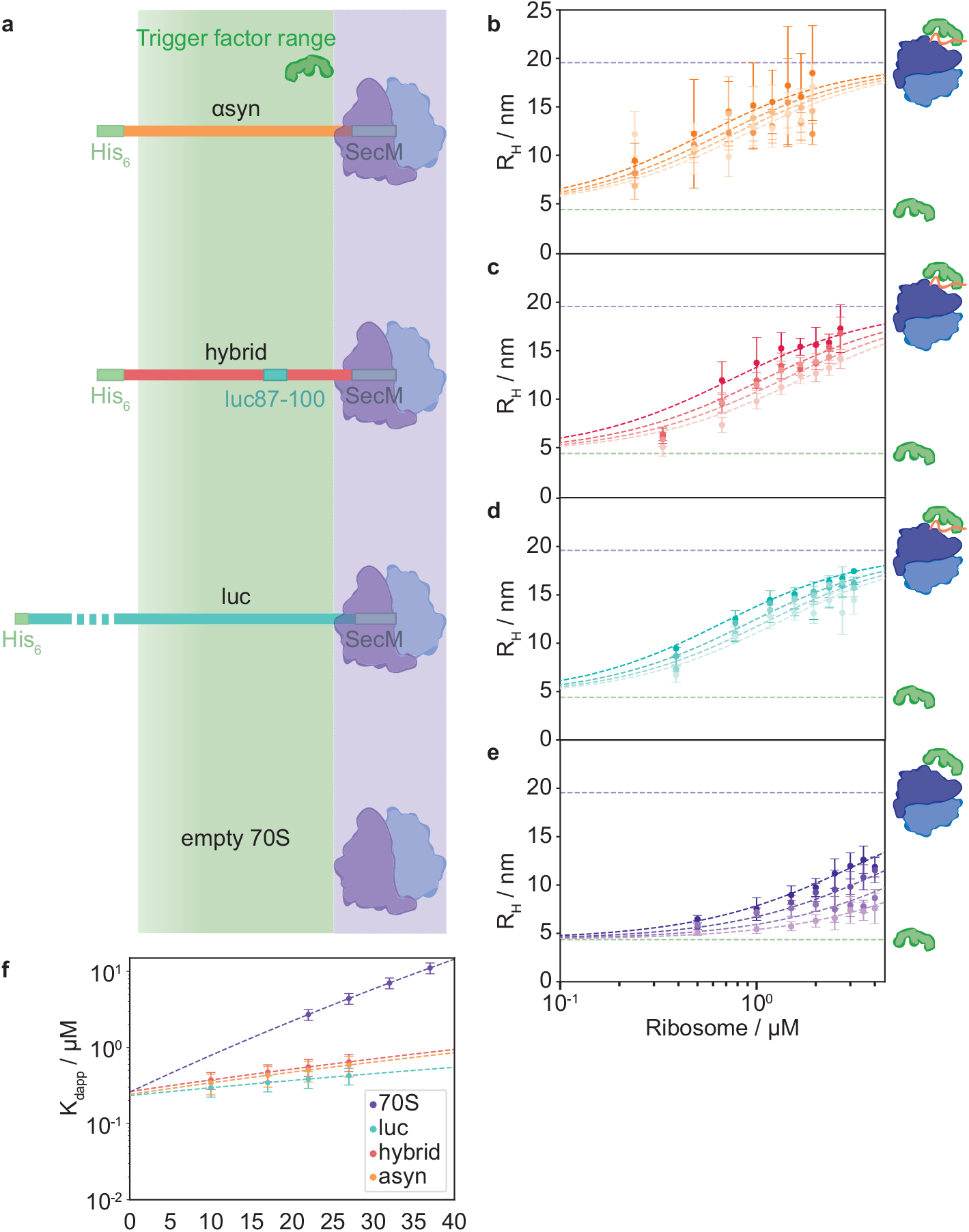
Nascent chain binding. **a** Schematic of the nascent chain constructs used in this study, comprising the SecM stall sequence, protein of interest, and a His_6_ tag for purification. The estimated range that can be contacted by the chaperone is shaded in green (residues 1-118).^19, 36^ The R_*H*_ of 200 nM TF as a function of RNC concentration and temperature (10–27^◦^C) for **b** *α*-synuclein, **c** *α*-synuclein(luc87-100), and **d** firefly luciferase. The estimated complex size from the intrinsic RNC fluorescence (grey dashed line) and *R*_*H*_ for TF (dashed green line) are shown. **e** TF binding to the empty 70S ribosome at 22–37^◦^C. **f** *K*_*d app*_ for global fits of ΔH and ΔS to the binding curves in **b - e**.

We developed a microfluidic platform for temperature-controlled binding measurements, which enables us to determine ΔH and ΔS for the interactions. We tested the setup by measuring the hydrodynamic radius for TF (*R*_*TF*_) for 10 - 37^◦^C, taking the changes in solution viscosity into account (1.31–0.69 mPa s for 10–37^◦^C), and we find that the size is constant with temperature (4.35 *±* 0.34 nm), SI Figure 2**b**.

We then acquired binding curves for TF and ribosome substrates at 10-37^◦^C, Figure 2**b-e**. ΔH and ΔS were used as free parameters in global fits to the binding data across four different temperatures (SI Table 1), using fixed values for the fractions of NC occupancy of the ribosomes (91% for *α*syn, 92% for the hybrid, and 40% for luc, determined by western blot, see Methods), *K*_*d*_ for dimerisation and binding to the empty ribosomes, see Methods, SI Figures 1**a** and SI Table 2. This analysis also yielded the ΔG and apparent *K*_*d*_ for the individual interactions (Figure 2**f** and SI Table 2). We found that TF binds to all three RNCs with nanomolar affinity (296–647 nM across 10–27^◦^C, 482*±*87 nM at 22^◦^C) and does not show a significant preference for luc, the NC we expected to be a particularly high-affinity substrate.^3, 7^ In agreement with literature reports, TF has a lower affinity for empty ribosomes, with *K*_*d*_ in the micromolar range (2.71*±*0.44–11.1*±*1.8 *μ*M for 22–37^◦^C).^3^ We thus see a step change in the chaperone affinity when the ribosome is occupied by an NC.

The N-terminal ribosome binding domain of TF is known to interact with the ribosomal protein uL23 when it docks at the exit tunnel of the 70S ribosome.^2, 13^ This interaction with the ribosome has been found to be necessary for NC engagement, and TF with a triple alanine mutation in the conserved ribosome binding sequence does not bind to RNCs.^7–9, 13^ In our snapshots of stalled RNCs we are observing the equilibrium for TF association with the RNC at the ribosome surface, rather than TF binding to the elongated NC. Our analysis of the thermodynamic driving forces behind TF function shows that the TF-70S interaction is driven by a negative ΔH and carries an entropy penalty (ΔH = -69.8 *±* 11.3 kJ mol^−1^ and ΔS = -132 *±* 21.5 J mol^−1 ◦^K^−1^, T*·*ΔS = 39.1 *±* 6.33 kJ mol^−1^ at 22^◦^C), Figure 3**a**.

**Figure 3:**
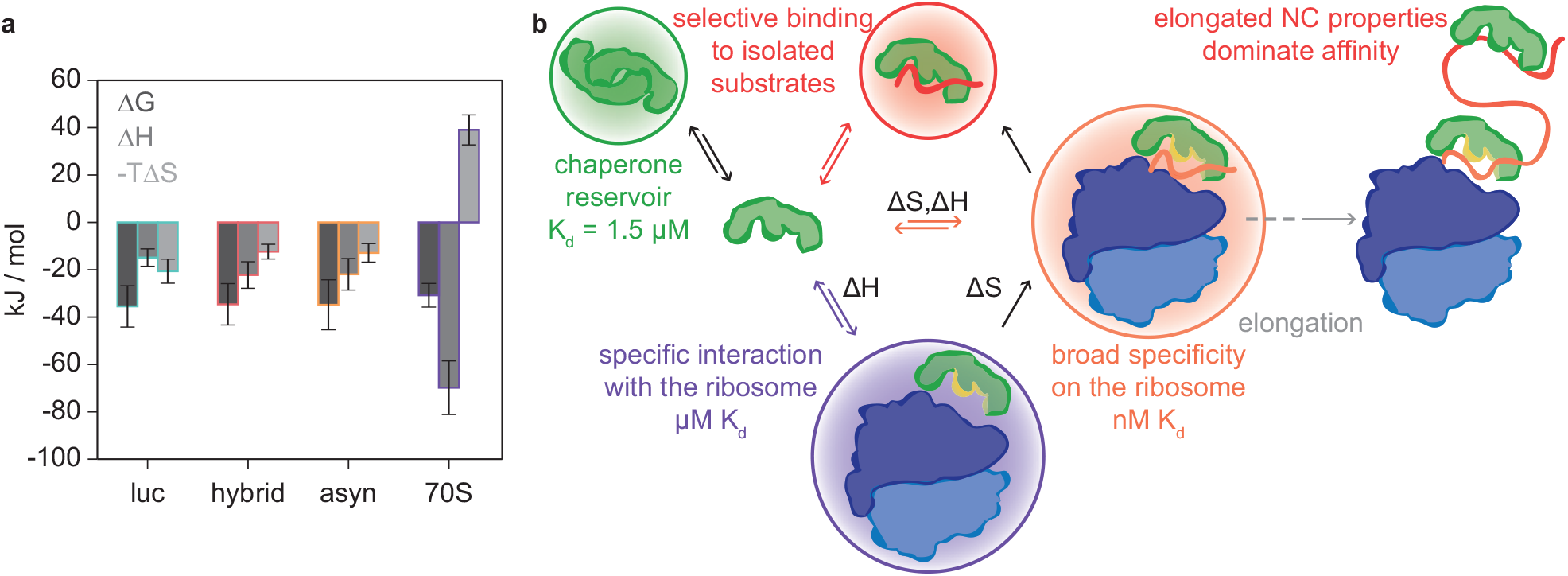
Thermodynamics of chaperone function. **a** The thermodynamic profiles for TF binding to three diverse RNC substrates show that binding is favoured by both ΔH and ΔS contributions, whereas TF binding to empty ribosomes is enthalpy-driven. **b** Summary schematic showing the thermodynamic driving forces and equilibrium parameters measured in this study in the context of TF function. The dimer conformation acts as a reservoir, which can release TF to compensate for a reduction in the monomer concentration. TF occupies a fraction of empty ribosomes due to the high total concentrations of TF and 70S (50 *μ*M and 30 *μ*M).^41, 42^ TF undergoes a conformational change upon binding to the ribosome (indicated by yellow surface), exposing hydrophobic patches and priming the chaperone for NC interactions.^3, 37, 38^ The similar thermodynamic profiles for the TF-RNC interactions suggest that TF employs a general strategy when associating with nascent chains at the ribosome surface, leading to similar *K*_*d*_ values. In contrast, when TF remains associated with an elongating nascent chain and leaves the ribosome surface during protein synthesis reports have found the duration of the complex to be substrate-dependent.^3^

The central question in this study is whether TF employs a general strategy for binding to its broad base of RNC substrates. Using global fits to the binding data for the three different RNCs, we found that the interactions show similar thermodynamic profiles, Figure 3**a**. The overall ΔΔG for TF binding to the RNCs versus empty ribosomes is -4–5 kJ mol^−1^ at 22^◦^C. The more negative ΔG for TF-RNC interactions is driven by a favourable entropy factor T*·*ΔS=12.4– 20.6 kJ mol^−1^, Figure 3**a**. This favourable Δ*S* is likely to arise from the release of ordered solvent at the TF substrate binding groove, which is exposed by a conformational change upon binding to the ribosome.^3, 37, 38^ These results suggest that TF employs two distinct thermodynamic strategies to achieve a favourable ΔG for the specific interaction with the ribosome via the TF ribosome binding domain (ΔH) and the general association with RNC substrates (ΔH and ΔS), Figure 3**b**.

We also investigated the difference in TF affinity for a potential substrate when encountered in isolation and as an RNC. IDPs are a particularly interesting case, because chaperone binding to the isolated protein would potentially deplete both the chaperone and IDP pool, thus inhibiting their native functions. Previous studies have shown limited interactions with *α*syn in free solution, cross-linking between TF and an *α*yn-RNC, but no specific interaction by fluorescence measurements on actively translating ribosomes.^7, 19, 39^ Although *α*syn is not in itself considered a ‘good’ TF substrate, the chaperone binds to the *α*syn RNC with higher affinity than to the empty ribosome (e.g. *K*_*dapp*_ = 482 *±* 87 nM and 2.71 *±*0.44 *μ*M at 22^◦^C.^19^ We did not detect binding between TF and isolated *α*syn ≤10 *μ*M, SI Figure 3. The increased affinity for the RNC is therefore not a simple combination of ribosome affinity and binding to an unfolded protein.

We consider the implications of the equilibrium parameters measured here for the distribution of TF between binding partners in the cell via two scenarios - *Escherichia coli* under slow and fast growth conditions, SI Figure 5. The fraction of actively translating ribosomes ranges from 20-90% between slow and fast growing *E. coli*,^40^ thus generating a large shift in the populations of high (RNCs) and low affinity (empty 70S) TF substrates. Both TF and ribosomes are present at relatively high concentrations in the cell, with TF at 50 *μ*M versus 30 *μ*M ribosomes.^41, 42^ We estimated the effect of this shift on the distribution of TF between states given the *K*_*d*_ values measured in this study, SI Figure 5, SI Table 2 Online Methods. Here, we investigate the explore the effect of ribosome occupancy, but this quantitative model can be used to explore the effect of other parameters (temperature, substrate affinity etc.), and it can be extended to include isolated substrates or additional pathways. When the concentration of high-affinity substrates (RNCs) increases from 6 to 27 *μ*M (slow to fast growth), the fractions of empty and active ribosomes occupied by TF remain similar at 62% and 90% (slow growth), and 59% and 89% (fast growth), while the TF monomer concentration remains steady (4.37 *μ*M and 3.90 *μ*M), thus favouring chaperone binding to high-affinity (*K*_*d*_ ≤ 1 *μ*M) substrates. The dimer population acts as the main source of TF for RNC-binding at 12.7 *μ*M for slow growth and 10.1 *μ*M for fast (equivalent to 20.2 *μ*M and 25.4 *μ*M in monomer). This balanced network of equilibria enables TF to provide flexible support for cellular proteostasis, Figure 3**b**.

## Discussion

We have employed quantitative microfluidic assays to measure TF-ligand binding directly in free solution, an approach that has allowed us to dissect the thermodynamic driving forces that underpin TFs role as the sole ribosome-associated chaperone in *E. coli*, Figure 3**b**. Here, we probe the interactions of TF in free solution, without the need for chemical cross-linking or downstream purification. The strategy, we have developed uses direct observations of the physicochemical properties of TF as reporters of binding. The equilibrium measurements of *K*_*d*_ = 1.5 *±* 0.25 *μ*M for TF dimerisation, *K*_*d*_ in the low *μ*M for ribosome binding (2.7 *±* 0.44 *μ*M), and *K*_*d*_s of 385 - 555 nM for RNC association confirm literature reports.^3, 4, 13, 16, 43^ In addition, our approach readily permits a dissection of the enthalpic and entropic contributions to the interactions and, as discussed below, shows how switching between enthalpic and entropic compensation governs the plethora of TF dynamic equilibria during translation. The method allows us to probe the subtlety of the parameters that modulate TF equilibria: a range of *K*_*d*_ values have been reported for central interactions depending on the experimental parameters, including dimerisation (*K*_*d*_ of 1–18 *μ*M),^3, 4, 13, 16^ and ribosome binding (e.g. 140 nM and ≈1 *μ*M).^18, 43^ The microfluidic techniques we use here, and in particular the microfluidic diffusional sizing assay, are flexible and allows for a comprehensive analysis to be carried out e.g. across different solution conditions, labelling strategies, and even in cell lysate.^30, 31^ We therefore envision that this platform will complement existing approaches in the biomolecular sciences, and that it can be used to resolve apparent inconsistencies in our understanding of TF function.

In our analysis of the thermodynamics of TF-substrate binding, we find that the most notable distinction is between TF binding to RNCs versus empty ribosomes, Figure 3**a**. Ribosome-association is driven by a negative ΔH = -69.8 *±* 11.3 kJ mol^−1^. This finding agrees with TF-70S binding being driven by specific interactions such as between GFRxGxxP sequence in the TF ribosome binding domain and complementary features on the ribosome surface e.g. Glu on L23.^2, 13^ The favourable ΔH compensates for an unfavourable −*T* ΔS = 39.1 *±*6.3 kJ mol^−1^, which is attributable to both the loss of conformational entropy upon complex formation and the loss of solvent entropy due to conformational changes exposing hydrophobic patches when TF docks at the ribosomal exit tunnel.^3, 37, 38^ Furthermore, once TF is associated with the ribosome, NC binding is effectively an intramolecular interaction, which benefits from not carrying the entropy penalty of complex formation and from a high effective concentration of TF at the exit tunnel. In addition, avidity effects due to additional interactions with the NC are likely to contribute to the increased affinity for RNCs. This configuration primes TF for interaction with a nascent polypeptide emerging from the exit tunnel. Indeed, we find that RNC binding for all three NCs has a favourable ΔS ranging from 41.9 *±* 10.6 to 69.9 *±* 17.2 J mol^−1 ◦^K^−1^, Figure 3**b**. The large TF substrate-binding surface is shielded at the centre of the TF dimer complex, Figure 3**b**.^29^ The conformational change upon TF binding to the ribosome^3, 37, 38^ may therefore be key to achieving high affinity for a broad substrate base without the need for energy input from ATP hydrolysis.

The ribosome surface is increasingly recognised as interacting with NCs and modulating their co-translational folding,^44^ these rapid interactions are not confined to the protection of hydrophobic residues, but can also be electrostatic.^45–47^ RNC-bound TF is in competition with the ribosome surface for binding to the NC, both types of interactions have been shown to inhibit folding.^1, 3, 46, 47^ The emergent polypeptide is then protected from misfolding and aberrant interactions by transient association with TF and the ribosome surface, providing a comprehensive system for shielding the newly synthesised polypeptide.

Our thermodynamic characterisation complements kinetic studies, where the *t*_1*/*2_s for TF docking on the ribosome were found to be similar for both empty and translating ribosomes (≈ 10 s).^3^ This step is required for further RNC interactions, and TF variants lacking the ribosome-binding motif do not compete with wild-type chaperone for RNC binding.^3, 7^ Regions of high hydrophobicity alone were therefore not sufficient for TF-NC association. However, TF can detach from the ribosome surface and remain associated with the growing NC, and the residence time depends on the properties of the NC, values for *t*_1*/*2_ of up to 35 or 110 s have been reported.^3, 7^ As the TF-NC complex moves away from the ribosome surface, the determinants for continued substrate-association approach those that govern selective TF binding to isolated substrates in solution, Figure 3**b**.^5, 27^ We found that the chaperone has a higher affinity for the *α*-synuclein-RNC than 70S alone (*K*_*d app*_ = 482 *±* 87 nM versus 2.71 *±* 0.44 *μ*M and *>*10 *μ*M at 22^◦^C), TF thus targets the *α*syn NC as a substrate, but is able to discriminate against IDP-binding in free solution, SI Figure 3.

We have investigated TF in equilibrium with RNCs that have been stalled mid-synthesis, reporting on TF interactions with the translating ribosome and nascent polypeptide. The chaperone continuously dissociates and re-associates with the ribosome surface regardless of whether a nascent chain is present.^3^ In our snapshots, we probe the apparent *K*_*d*_ and thermodynamic parameters for TF-RNC binding at the ribosomal exit tunnel, and we find that the initial TF-RNC interaction has a low dependence on the NC properties, e.g. the presence of hydrophobic target motifs such as in the hybrid RNC. These results suggest that TF associates to a high degree with newly synthesised peptides as they emerge from the ribosomal exit tunnel, and that the chaperone achieves nanomolar RNC affinity through a common strategy for diverse NCs. The variation in *t*_1*/*2_ between different NCs on actively translating ribosomes indicates that selective support is then provided to nascent chains with e.g. stretches of hydrophobic residues as they grow.^3^

We have developed a robust and easy to use experimental strategy with potential applications in both fundamental and translational research in the biomolecular sciences and medicine. Unlike the selectivity displayed by TF for isolated proteins in solution, we find that the chaperone interacts with the very diverse set of RNCs with similar (nM) affinity, mediated by a favourable ΔS. Taken together, our results suggest a general strategy for RNC association, which does not rely on specific sequence properties in the NC. TF elegantly combines high-affinity RNC binding to achieve its ubiquitous function with substrate-dependent kinetics of dissociation from the elongating NC, offering extended protection to selected NCs. These observations reconcile the two roles of TF as selective when engaging isolated substrates and the exceptionally broad function of TF as a co-translational chaperone.

## Data availability

Source data for figures is available online. Microfluidic image analysis was performed as outlined in Herling et al. for electrophoresis,^30^ code for the microfludic diffusional sizing analysis is available at https://zenodo.org/record/3881940#.Y-TgAxPP0bZ.^31^ Additional data available upon reasonable request.

## Acknowledgements

We would like to thank the BBSRC (T.W.H., T.P.J.K.), Oppenheimer Foundation (T.W.H.), Murray Edwards College Cambridge (T.W.H.), Newman Foundation (T.P.J.K.), and the Wellcome Trust (A.C., L.C., J.C.) for funding this research. We would like to thank Prof. F.-Ulrich Hartl (Max Planck Institute of Biochemistry) for the TF construct.

## Online Methods

### Microfluidic device preparation

Microfluidic devices were cast in PDMS (Momentive RTV615, Techsill, UK) using standard soft lithography methods^48^. The clear PDMS was coloured black by the addition of a small quantity of carbon nanopowder, 0.2% w/w, prior to curing (Sigma, UK). Inlet and outlet holes were punched using a biopsy punch (WPI, Florida, US). The PDMS devices were bonded to glass slides in a plasma oven using an oxygen plasma (Diener Electronics, Germany). The electrodes were fabricated by placing the bonded device glass slide down on a hot plate set to 79^◦^ C and inserting InBiSn alloy (51% In, 32.5% Bi, 16.5% Sn, Conro Electronics, UK) through the solder inlet.^30^

### Microfluidic measurements

In order to follow binding equilibria as a function of temperature, we have enhanced a custom built microscope with a temperature-controlled stage equipped with Peltier elements and a proportional integral derivative (PID) controller, enabling us to both heat and cool the sample on chip and paving the way for a full thermodynamic analysis, Figure 1 and SI Figure 2**a**.^35^ To detect multiple components in a mixture, we combine selective fluorophore labelling of trigger factor with AlexaFluor488 with the intrinsic fluorescence of aromatic amino acids such as tryptophan and tyrosine, Figure 1**c-d** and SI Figure 4.^34^ Samples were incubated at the relevant temperature for 30 min prior to loading on chip.

### Sample preparation

Trigger factor N326C was expressed and purified as previously described.^19^ The chaperone was selectively labelled with AlexaFluor488 maleimide according to the manufacturers instructions (ThermoFisher Scientific) at position 326, a labelling position used in literature reports.^3, 7^

Ribosomes and RNCs were prepared and purified as previously reported, see SI for construct sequences.^36^ RNC occupancy was assessed by western blot.^36^ The fraction of ribosomes occupied by an NC (*α*) was high when expression was performed in cells containing native trigger factor: 91% for *α*-synuclein; 92% for the hybrid construct;. Both synuclein-based RNCs were purified with only small amounts of trigger factor present in the final sample, 1% and 2% of RNC concentration for *α*-Synuclein and the hybrid construct respectively. However, a considerable proportion of the luciferase RNC co-purified with trigger factor at 22% of the RNC concentration, suggesting that trigger factor has a high affinity for this substrate. We therefore purified the firefly luciferase RNC for the microfluidic measurements in a TF knockout strain, E. coli Δtig, with a final occupancy of 40%.^36^ The concentration of unlabelled trigger factor was taken into account in the data analysis, se SI.

### Binding curve analysis

*TF*_*t*_ was the monomeric TF (*m*), dimeric TF (*d*), TF on empty ribosomes (*TF* _70*S*_) and RNCs (*TF*_*RNC*_):

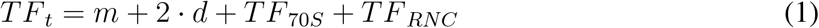

We used three *K*_*d*_s for TF complex formation to express *d, TF* _70*S*_ and *TF*_*RNC*_ in terms of of *m* and the total 70S/RNC concentration (*r*_0_),

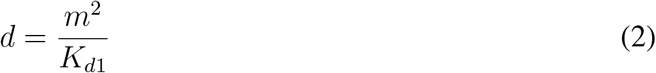

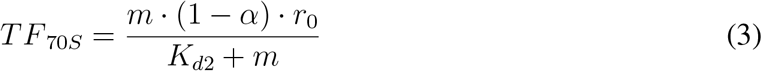

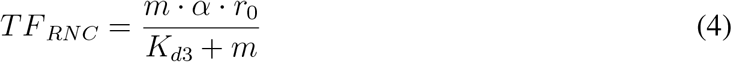

The observed hydrodynamic radius (*R*_*obs*_) for the total TF depended on the contributions from isolated TF (*R*_*TF*_) and 70S or RNC-bound TF (*R*_*complex*_):

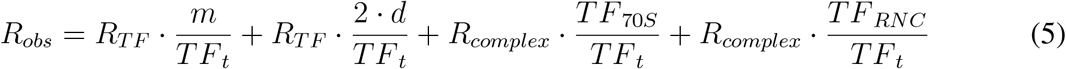

In this analysis, we did not distinguish the the monomer/dimer radii, as *R*_*TF*_ varied little above and below *K*_*D*1_ in agreement with literature reports, see SI Figure 3**b**.^28^ *R*_*TF*_ was fixed to that of 200 nM TF (4.35 *±* 0.34 nm) and *R*_*complex*_ was set to the average value measured by intrinsic fluorescence for the 70S and RNC samples (19.6 *±* 1.4 nm), SI Figure 3**a**.

As the changes to *R*_*TF*_ upon dimerisation are small, SI Figure 3**b**,^28^ we measured *K*_*d*1_ by observing the electrophoretic mobility as a function of TF concentration, SI Figure 2.^30, 49^

The *K*_*d*_ for binding to the 70S ribosome and RNCs was determined by a global fit of Δ*H* and Δ*S* to binding curves acquired at four different temperatures using the van’t Hoff equation, Figure 2,

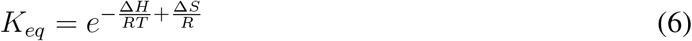

with *K*_*d*_ = 1*/K*_*eq*_. We first determined Δ*H* and Δ*S* for binding to empty ribosomes and used these values to determine *K*_*d*2_ when fitting the RNC data. The errors on fit parameters were determined by combining the relative errors from *R*_*TF*_, *R*_*complex*_, residuals for the fit, *R*_*Hobs*_ (between triplicates), and an estimate of 20% on the RNC occupancy.

We used Equations 1-4 to calculate the TF distributions between states for slow and fast growing cells.^40^ The calculations were made for *K*_*d*_ values at 22^◦^C to avoid extrapolating from the measured temperature range. Cellular ribosome and TF concentrations were estimated as 30 *μ*M and 50 *μ*M respectively for the purpose of this illustration.^41, 42^ As an approximation for the RNC affinity, we use the average *K*_*d*_ for RNC binding (here, 451 nM).

## References

1. Agashe, V. R. et al. Function of Trigger Factor and DnaK in Multidomain Protein Folding: Increase in Yield at the Expense of Folding Speed. Cell 117, 199–209 (2004).

2. Ferbitz, L. et al. Trigger factor in complex with the ribosome forms a molecular cradle for nascent proteins. Nature 431, 9396–9401 (2004).

3. Kaiser, C. M. et al. Real-time observation of trigger factor function on translating ribosomes. Nature 444, 455–460 (2006).

4. Rutkowska, A. et al. Dynamics of Trigger Factor Interaction with Translating Ribosomes. Journal of Biological Chemistry 283, 4124–4132 (2008).

5. Martinez-Hackert, E. & Hendrickson, W. A. Promiscuous Substrate Recognition in Folding and Assembly Activities of the Trigger Factor Chaperone. Cell 138, 923–934 (2009).

6. Hoffmann, A., Bukau, B. & Kramer, G. Structure and function of the molecular chaperone Trigger Factor. Biochimica et Biophysica Acta 1803, 650–661 (2010).

7. Lakshmipathy, S. K., Gupta, R., Pinkert, S., Etchells, S. A. & Hartl, F. U. Versatility of trigger factor interactions with ribosome-nascent chain complexes. Journal of Biological Chemistry 285, 27911–27923 (2010).

8. Oh, E. et al. Selective Ribosome Profiling Reveals the Cotranslational Chaperone Action of Trigger Factor In Vivo. Cell 147, 1295–1308 (2011).

9. Hoffmann, A., Becker, A. H., Zachmann-brand, B., Deuerling, E. & Bukau, B. Concerted Action of the Ribosome and the Associated Chaperone Trigger Factor Confines Nascent Polypeptide Folding. Molecular Cell 48, 63–74 (2012).

10. Mashaghi, A. et al. Reshaping of the conformational search of a protein by the chaperone trigger factor. Nature 500, 98–101 (2013).

11. Zhao, L. et al. Bacterial RF3 senses chaperone function in co-translational folding. Molecular Cell 81, 2914–2928.e7 (2021).

12. Deuerling, E., Schulze-specking, A., Tomoyasu, T., Mogk, A. & Bukau, B. Trigger factor and DnaK cooperate in folding of newly synthesized proteins. Nature 133, 693–696 (1999).

13. Kramer, G. et al. L23 protein functions as a chaperone docking site on the ribosome. Nature 419, 171–174 (2002).

14. Merz, F. et al. Molecular mechanism and structure of Trigger Factor bound to the translating ribosome. The EMBO Journal 27, 1622–1632 (2008).

15. Nilsson, O. B., Muller-Lucks, A., Kramer, G., Bukau, B. & Von Heijne, G. Trigger Factor Reduces the Force Exerted on the Nascent Chain by a Cotranslationally Folding Protein. Journal of Molecular Biology 428, 1356–1364 (2016).

16. Patzelt, H. et al. Three-state equilibrium of escherichia coli trigger factor. Biological Chemistry 383, 1611–1619 (2002).

17. Saio, T., Guan, X., Rossi, P., Economou, A. & Kalodimos, C. G. Structural Basis for Protein Antiaggregation Activity of the Trigger Factor Chaperone. Science 344, 1250494 (2014).

18. Bornemann, T., Holtkamp, W. & Wintermeyer, W. Interplay between trigger factor and other protein biogenesis factors on the ribosome. Nature Communications 5 (2014).

19. Deckert, A. et al. Structural characterization of the interaction of α-synuclein nascent chains with the ribosomal surface and trigger factor. Proceedings of the National Academy of Sciences 113, 5012–5017 (2016).

20. Yang, F. et al. Single-molecule dynamics of the molecular chaperone trigger factor in living cells. Molecular Microbiology 102, 992–1003 (2016).

21. Haldar, S., Tapia-Rojo, R., Eckels, E. C., Valle-Orero, J. & Fernandez, J. M. Trigger factor chaperone acts as a mechanical foldase. Nature Communications 8, 668 (2017).

22. Kawagoe, S., Nakagawa, H., Kumeta, H., Ishimori, K. & Saio, T. Structural insight into proline cis/trans isomerization of unfolded proteins catalyzed by the trigger factor chaperone. The Journal of Biological Chemistry 293, 15095–15106 (2018).

23. Wu, K., Minshull, T. C., Radford, S. E., Calabrese, A. N. & Bardwell, J. C. Trigger factor both holds and folds its client proteins. Nature Communications 13, 1–15 (2022).

24. De Geyter, J. et al. Trigger factor is a bona fide secretory pathway chaperone that interacts with SecB and the translocase. EMBO Reports 21, 1–17 (2020).

25. Rizzolo, K. et al. Functional cooperativity between the trigger factor chaperone and the ClpXP proteolytic complex. Nature Communications 12, 281 (2021).

26. Saio, T., Kawagoe, S., Ishimori, K. & Kalodimos, C. G. Oligomerization of a molecular chaperone modulates its activity. eLife 7, e35731 (2018).

27. Mitra, R., Wu, K., Lee, C. & Bardwell, J. C. A. ATP-Independent Chaperones. Annual Review of Biophysics 51, 409–429 (2022).

28. Rathore, Y. S., Dhoke, R. R., Badmalia, M. & Sagar, A. SAXS Data Based Global Shape Analysis of Trigger Factor (TF) Proteins from E. coli, V. cholerae, and P. f rigidicola: Resolving the Debate on the Nature of Monomeric and Dimeric Forms. Journal of Physical Chemistry B 119, 6101–6112 (2015).

29. Morgado, L., Burmann, B. M., Sharpe, T., Mazur, A. & Hiller, S. The dynamic dimer structure of the chaperone Trigger Factor. Nature Communications 8, 1992 (2017).

30. Herling, T. W. et al. Integration and characterization of solid wall electrodes in microfluidic devices fabricated in a single photolithography step. Applied Physics Letters 102 (2013).

31. Arosio, P. et al. Microfluidic diffusion analysis of the sizes and interactions of proteins under native solution conditions. ACS Nano 10, 333–341 (2016).

32. Zhang, Y. et al. A microfluidic strategy for the detection of membrane protein interactions. Lab on a Chip 20, 3230–3238 (2020).

33. Schneider, M. M. et al. The Hsc70 disaggregation machinery removes monomer units directly from α -synuclein fibril ends. Nature Communications 12, 1–11 (2021).

34. Challa, P. K. et al. Real-Time Intrinsic Fluorescence Visualization and Sizing of Proteins and Protein Complexes in Microfluidic Devices. Analytical Chemistry 90, 3849–3855 (2018).

35. Feng, Y. & Wandinger-ness, A. Affordable controlled-temperature microscope slide for live cell viewing. Technical Tips Online 3, 83–87 (1998).

36. Cassaignau, A. M. E. et al. A strategy for co-translational folding studies of ribosome-bound nascent chain complexes using NMR spectroscopy. Nature Protocols 11, 1492–1507 (2016).

37. Baram, D. et al. Structure of trigger factor binding domain in biologically homologous complex with eubacterial ribosome reveals its chaperone action. Proceedings of the National Academy of Sciences 102, 12017–12022 (2005).

38. Deeng, J. et al. Dynamic Behavior of Trigger Factor on the Ribosome. Journal of Molecular Biology 428, 3588–3602 (2016).

39. Tomic, S., Johnson, A. E., Hartl, F. U. & Etchells, S. A. Exploring the capacity of trigger factor to function as a shield for ribosome bound polypeptide chains. FEBS Letters 580, 72–76 (2006).

40. Dai, X. et al. Reduction of translating ribosomes enables Escherichia coli to maintain elongation rates during slow growth. Nature Microbiology 2, 16231 (2016).

41. Lill, R., Crooke, E., Guthrie, B. & Wickner, W. The Trigger Factor CycE Includes Ribosomes, Presecretory Proteins, and the Plasma Membrane. Cell 54, 1013–1018 (1988).

42. Bremer, H. & Dennis, P. Escherichia coli and Salmonella: cellular and molecular biology (ASM Press, Washington DC, 1996).

43. Maier, R., Eckert, B., Scholz, C., Lilie, H. & Schmid, F. X. Interaction of trigger factor with the ribosome. Journal of Molecular Biology 326, 585–592 (2003).

44. Alexander, L. M., Goldman, D. H., Wee, L. M. & Bustamante, C. Non-equilibrium dynamics of a nascent polypeptide during translation suppress its misfolding. Nature Communications 10, 1–11 (2019).

45. Plessa, E. et al. Nascent chains can form co-translational folding intermediates that promote post-translational folding outcomes in a disease-causing protein. Nature Communications 12 (2021).

46. Cassaignau, A. M. E. et al. Interactions between nascent proteins and the ribosome surface inhibit co-translational folding. Nature Chemistry 13, 1214–1220 (2021).

47. Chan, S. H. S. et al. The ribosome stabilizes partially folded intermediates of a nascent multidomain protein. Nature Chemistry 14, 1165–1173 (2022).

48. McDonald, J. C. & Whitesides, G. M. Poly (dimethylsiloxane) as a material for fabricating microfluidic devices. Accounts of Chemical Research 35, 491–499 (2002).

49. Herling, T. W. et al. A Microfluidic Platform for Real-Time Detection and Quantification of Protein-Ligand Interactions. Biophysical Journal 110, 1957–1966 (2016).

