## Supplementary Information for "Thermodynamic profiles for co-translational trigger factor function"

### Supplementary Information: Thermodynamic profiles for co-translational trigger factor function

**Protein constructs and sequences**

RNC constructs were prepared and purified as previously reported.[1] The constructs contain an N-terminal hexa-His tag and a C-terminal SecM sequence.

**Luciferase RNC**

HHHHHHASME[10] DAKNIKKGPA[20] PFYPLEDGTA[30] GEQLHKAMKR[40] YALVPGTIAF[50]

TDAHIEVNIT[60] YAEYFEMSVR[70] LAEAMKRYGL [80] NTNHRIVVC[90] SENSLQFFMPV[100]

LGALFIGVAV[110] APANDIYNER[120] ELLNSMNISQ[130] PTVVFVSKKG[140] LQKILNVQKK[50]

LPIIQKIIIM[160] DSKTDYQGFQ[170] SMYTFVTSHL[180] PPGFNEYDFV[190] PESFDRDKTI[200]

ALIMNSSGST$^{210}$ GLPKGVALPH$^{220}$ RTACVRFSHA$^{230}$ RDPIFGNQII$^{240}$ PDTAILSVVP$^{250}$

FHHGFGMFTT$^{260}$ LGYLICGFRVV$^{270}$ LMYRFEEEL$^{280}$ FLRSLQDYKI$^{290}$ QSALLVPTLF$^{300}$

SFFAKSTLID$^{310}$ KYDLSNLHEI$^{320}$ ASGGAPLSKE$^{330}$ VGEAVAKRFH$^{340}$ LPGIRQGYGL$^{350}$

TETTSAILIT$^{360}$ PEGDDKPGAV$^{370}$ GKVVPFFEAK$^{380}$ VVDLDTGKTL$^{390}$ GVNQRGELCV$^{400}$

RGPMIMSGYV$^{410}$ NNPEATNALI$^{420}$ DKDGWLHSGD$^{430}$ IAYWDEDEHF$^{440}$ FIVDRLKSLI$^{450}$

KYKGYQVAPA$^{460}$ ELESILLQHP$^{470}$ NIFDAGVAGL$^{480}$ PDDDAGELPA$^{490}$ AVVVLEHGKT$^{500}$

MTEKEIVDYV$^{510}$ ASQVTTAKKL$^{520}$ RGGVVFVDEV$^{530}$ PKGLTGKLDA$^{540}$ RKIREILIKA$^{550}$

KKGGKSKLTS$^{560}$ EFFSTPVWIW$^{570}$ WWPRIRGPPP$^{580}$ PWT

**Hybrid RNC**

MHHHHHHASM$^{10}$ DVFMKGLSKA$^{20}$ KEGVVAAAEK$^{30}$ TKQGVAEAAG$^{40}$ KTKEGVLYVG$^{50}$

SKTKEGVVHG$^{60}$ VATVAEKTKE$^{70}$ QVTNVGGAVV$^{80}$ TGVTAVAQKT$^{90}$ VEGAG**QFFMP**$^{100}$

**VLGALFIGV**G$^{110}$ KNEEGAPQEG$^{120}$ ILEDMPVDPD$^{130}$ NEAYEMPSEE$^{140}$ GYQDYEPEAG$^{150}$

TTSEFFSTPV$^{160}$ WIWWWPRIRG$^{170}$ PPPPWT

The 13 amino acid insert from firefly luciferase (87-100) is shown in bold.

**$\alpha$-Synuclein RNC**

MHHHHHHENL$^{10}$ YFQGASMDVF$^{20}$ MKGLSKAKEG$^{30}$ VVAAAEKTKQ$^{40}$ GVAEAAGKTK$^{50}$

EGVLYVGSKT$^{60}$ KEGVVHGVAT$^{70}$ VAEKTKEQVT$^{80}$ NVGGAVVTGV$^{90}$ TAVAQKTVEG$^{100}$

AGSIAAATGF$^{110}$ VKKDQLGKNE$^{120}$ EGAPQEGILE$^{130}$ DMPVDPDNEA$^{140}$ YEMPSEEGYQ$^{150}$

DYEPEAGTTS$^{160}$ EFFSTPVWIW$^{170}$ WWPRIRGPPP$^{180}$ PWT

The isolated $\alpha$syn protein has the His-tag, but not the SecM sequence.

Table S1: Free energy contributions from global fits to the binding curves in Figure 2

| Ligand | $\Delta$H / kJ mol$^{-1}$ | $\Delta$S / J mol$^{-1}$ $^{\circ}$K$^{-1}$ |
|---|---|---|
| Luciferase | -14.9$\pm$3.66 | 69.9$\pm$17.2 |
| Hybrid | -22.2$\pm$5.61 | 41.9$\pm$10.6 |
| $\alpha$-synuclein | -21.9$\pm$6.65 | 43.7$\pm$13.2 |
| 70S ribosome | -69.8$\pm$11.3 | -132$\pm$21.5 |

Table S2: $K_{d\,app}$ calculated from the free energy contributions in SI Table 1

| Temperature / $^{\circ}$C | 10 | 17 | 22 | 27 | 32 | 37 | Est. error / % |
|---|---|---|---|---|---|---|---|
| Luciferase | 296 nM | 346 nM | 385 nM | 427 nM | – | – | 24.6 |
| Hybrid | 375 nM | 473 nM | 555 nM | 647 nM | – | – | 25.2 |
| $\alpha$-synuclein | 343 nM | 432 nM | 505 nM | 588 nM | – | – | 30.3 |
| 70S ribosome | – | – | 2.71 $\mu$M | 4.41 $\mu$M | 7.04 $\mu$M | 11.1 $\mu$M | 16.2 |

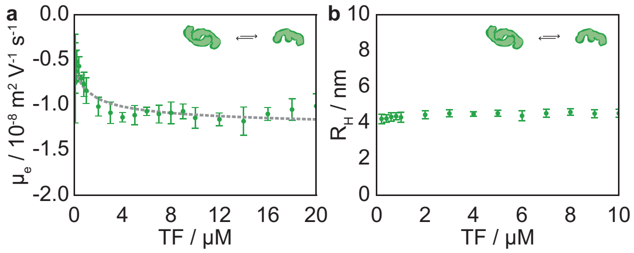


Figure S1: **a** Trigger factor self-associates to form a dimer. The average electrophoretic mobility is monitored as a function of trigger factor concentration. The dashed line shows a fit to of the data to give a $K_d$ of 1.5 $\mu$M, following the electrophoretic mobility from $\mu_{\text{monomer}} = -0.5 \cdot 10^{-8}\text{m}^2\,\text{V}^{-1}\,\text{s}^{-1}$ to $\mu_{\text{dimer}} = -1.3 \cdot 10^{-8}\text{m}^2\,\text{V}^{-1}\,\text{s}^{-1}$. Error bars show the standard deviation for three independent measurements. **b** The hydrodymanic radius of trigger factor as a function of concentration.

1. Deckert, A. *et al.* Structural characterization of the interaction of $\alpha$-synuclein nascent chains with the ribosomal surface and trigger factor. *Proceedings of the National Academy of Sciences* **113**, 5012–5017 (2016).

2. Lill, R., Crooke, E., Guthrie, B. & Wickner, W. The Trigger Factor CycE Includes Ribosomes , Presecretory Proteins , and the Plasma Membrane. *Cell* **54**, 1013–1018 (1988).

3. Bremer, H. & Dennis, P. *Escherichia coli and Salmonella: cellular and molecular biology* (ASM Press, Washington DC, 1996).

4. Kaiser, C. M. *et al.* Real-time observation of trigger factor function on translating ribosomes. *Nature* **444**, 455–460 (2006).

5. Dai, X. *et al.* Reduction of translating ribosomes enables Escherichia coli to maintain elongation rates during slow growth. *Nature Microbiology* **2**, 16231 (2016).

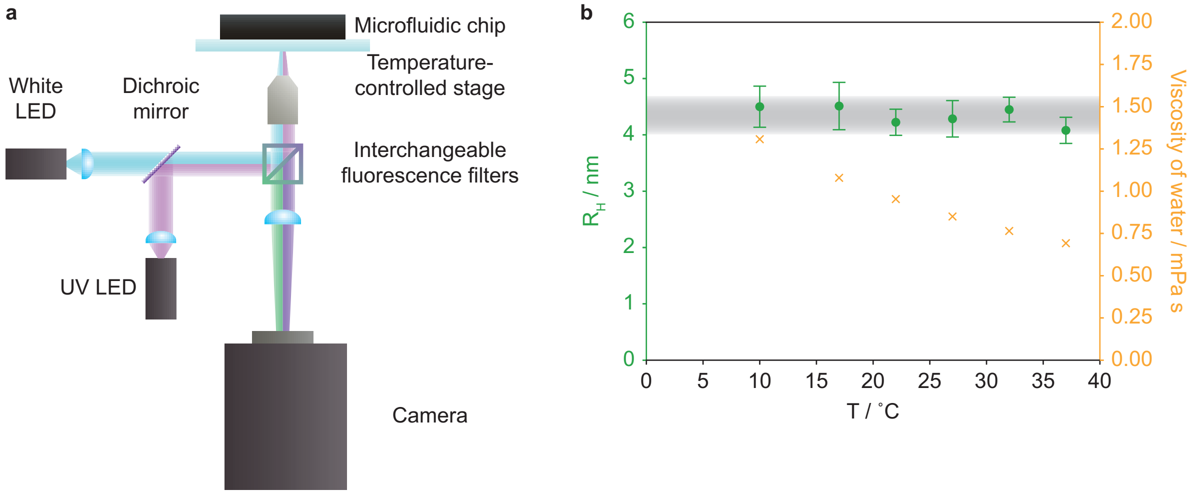


Figure S2: **a** Custom built microscope, the sample is illuminated with a white or a UV LED at 285 nm, fluorescence measurements are made using interchangeable filter sets to select the wavelength of interest. **b** $R_{TF}$ measured as a function of temperature is constant (green). Orange x shows the viscosity of water, which almost halves across the temperature range investigated here.

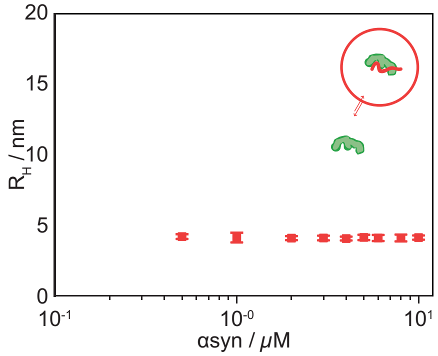


Figure S3: Trigger factor binding to potential ligands is monitored via microfluidic diffusional sizing. The $\mathrm{R_H}$ of 200 nM AlexaFluor488 labelled trigger factor is measured as a function of ligand concentration for isolated $\alpha$-Synuclein (red)

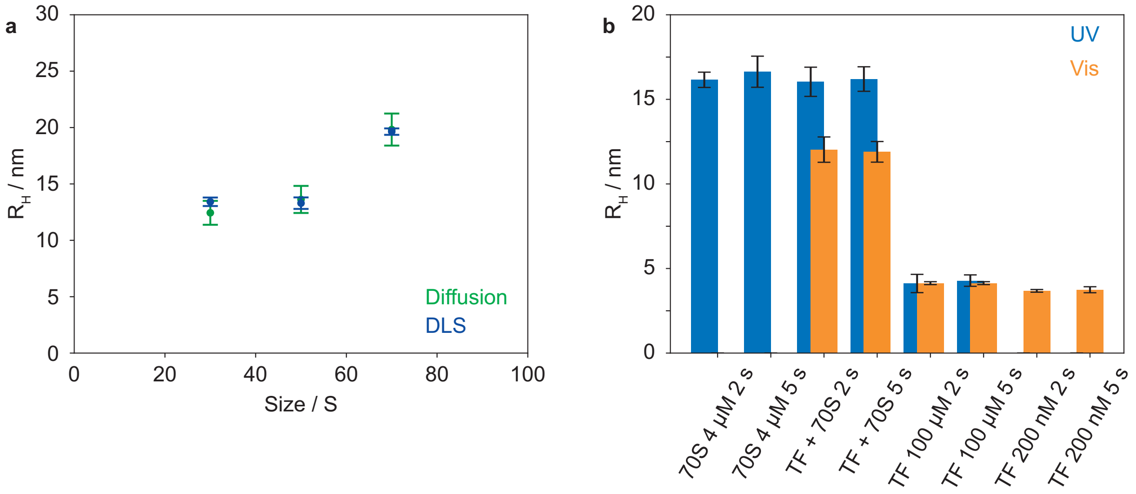


Figure S4: **a** $R_H$ for intact 70S ribosomes, the large subunit (50S) and small subunit (30S) measured by microfluidic diffusional sizing (2 $\mu$M) and dynamic light scattering (DLS). Error bars represent the standard deviation for three independent measurements. **b** Varying the exposure time (2 s and 5 s) in microfluidic diffusional sizing of: 4 $\mu$M 70S; 4 $\mu$M 70S + 200 nM TF; 100 $\mu$M TF sized through intrinsic fluorescence and fluorophore label; and 200 nM TF. Error bars are standard deviation for three independent measurements, except 4 $\mu$M 70S where two measurements were taken.

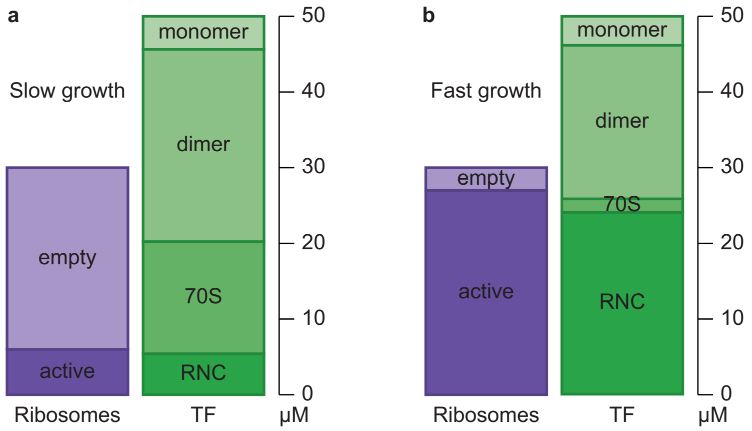


Figure S5: **a** TF and ribosome distribution in slow and **b** fast growing cells based on the $K_d$s measured here at 22°C ($K_d = 1.5$ $\mu$M for TF dimerisation; average $K_{d\,app} = 482$ nM for RNC binding; $K_d = 2.7$ $\mu$M for binding to empty ribosomes).[2–5] The total TF and ribosome concentrations used were 50 $\mu$M and 30 $\mu$M respectively.[2,3] The equilibria would be shifted by TF binding to isolated proteins.
